# Nunchaku: Optimally partitioning data into piece-wise linear segments

**DOI:** 10.1101/2023.05.26.542406

**Authors:** Yu Huo, Hongpei Li, Xiao Wang, Xiaochen Du, Peter S. Swain

## Abstract

When analysing two-dimensional data sets, scientists are often interested in regions where one variable depends linearly on the other. Typically they use an *ad hoc* method to do so. Here we develop a statistically rigorous, Bayesian approach to infer the optimal partitioning of a data set into contiguous piece-wise linear segments. Our `nunchaku` algorithm is freely available. Focusing on microbial growth, we use `nunchaku` to identify the range of optical density where the density is linearly proportional to the number of cells and to automatically find the regions of exponential growth for both *Escherichia coli* and *Saccharomyces cerevisiae*. For budding yeast, we consequently are able to infer the Monod constant for growth on fructose. Our algorithm lends itself to automation and high throughput studies, increases reproducibility, and will facilitate data analysis for a broad range of scientists.

## Introduction

A common scientific problem is understanding the relationship between two variables. When the dependent variable, or some transformation of it, depends linearly on the independent variable, the underlying system linking the two often behaves more simply than generally. As a consequence, scientists commonly focus their efforts on identifying and understanding this linear regime.

A well-known example is the growth of a population of cells. In log phase, when the logarithm of the number of cells increases linearly with time, the total mass of every intracellular component is growing exponentially and the mass per cell is approximately constant. Such steady-state conditions regularise growth, metabolic fluxes are balanced, and physiology simplifies, generating behaviours controlled by only a handful of variables [1].

Biologists therefore often wish to determine when growth is in log phase. Historically the approach has been to plot the logarithm of a variable correlating with the number of cells, such as optical density (OD), against time and to identify a linear region by eye [2]. Today this subjective technique is still used, with one scientist’s linear region not necessarily the same as another’s.

A challenge to developing alternative objective approaches is identifying a suitable non-linear model with which to compare the linear one. There is no general way to describe all relationships that we may observe. With a mechanistic understanding, we might generate a non-linear description, but such an understanding is often lacking and, anyhow, may obviate the need to find linear regimes.

Here we circumvent this problem by inferring the piece-wise linear description that best approximates an entire bivariate data set. We use a Bayesian approach and compare the evidence for every possible piece-wise linear description, found by marginalising over all possible linear fits to all possible contiguous subdivisions of the data. For the optimal choice, we provide statistics for each linear segment, allowing users to select straightforwardly the segment or segments of most interest. We implement our algorithm in Python and it is freely available. To illustrate its use, we discuss two examples: determining the range of OD of a liquid culture where the OD depends linearly on the number of cells and finding the exponential phases of microbial growth curves.

## Results

### Approximating data with a piece-wise linear model

Although our goal is to allow scientists to choose objectively the segment of their data that is ‘most’ linear, we adopt a general approach. For a two-dimensional data set, we will infer its optimal piece-wise linear description — the number of contiguous segments into which we should divide the data, where the boundaries of each of those segments should be, and the best-fit line for each segment. Deciding which of these segments is then best for the task in hand is unavoidably subjective. It is straightforward, however, to compare different segments by comparing properties of their best-fit lines, such as their gradients or *R*^2^ value — how much of the variance of the dependent variable is explained by the independent one [3].

We use a Bayesian approach to infer the best piece-wise linear model and assume only that the data of each segment is normally distributed around a line (Materials & Methods). To proceed analytically we marginalise over all gradients and *y*-intercepts of the lines for each segment using a mild approximation and choose the optimal number of segments by comparing marginal likelihoods. The data points bounding each segment are then estimated by the means of their posterior distribution. We consider the case with known measurement error separately from an unknown one and call our algorithm `nunchaku`.

### Verifying our approach

To verify our methodology (Materials & Methods), we generated synthetic data using piece-wise linear functions where we know the number of segments and their gradients, added Gaussian noise, and then inferred from this data the optimal number of segments and the gradients of the best-fit lines, assuming that we know magnitude of the measurement noise (Fig. 1A).

**Figure 1.**
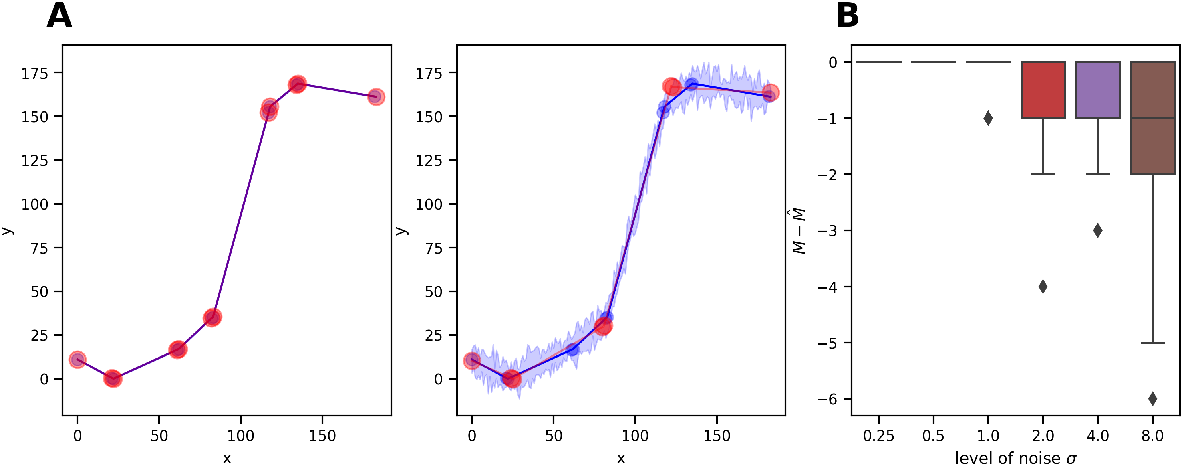
The `nunchaku` algorithm correctly predicts the number of linear segments in synthetic data providing the measurement noise is not too high. **(A)** Examples of 10 synthetic data sets with the ground truth in blue and twice the standard deviation shaded. The red circles are the predicted boundaries of each linear segment with the best-fit line in red. Left: with a measurement error of 0.25, the predictions overlap the data; Right: with a measurement error of 0.8, the predictions miss some segments which the noise obscures. As a prior, we specify only that the gradient of each line lies between [−25, 25]. **(B)** The algorithm underestimates the number of linear segments only once the magnitude of the measurement noise becomes sufficiently high. The actual number of segments is *M*; the estimated number is 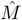.

The algorithm predicts correctly the correct of segments when the noise in the data is sufficiently low (Fig. 1B), but underestimates this number when the noise is larger. Such noise tends to blur two neighbouring segments so they seem one, rather than cause a single segment to appear as two or more. Similarly, if we decrease the angle between neighbouring segments, the noise is more likely to make two neighbouring segments appear contiguous, and the algorithm’s accuracy falls (Fig. S1B).

We confirmed that when the algorithm predicts correctly the number of segments, it also predicts correctly the gradient of the lines generating the data in the segments (Fig. S1C). As expected, this accuracy falls too with more noisy data.

When the measurement error is unknown, the results are similar (Fig. S3), but the algorithm runs slower because we must numerically integrate over all possible magnitudes of the measurement noise.

### Application 1: Finding the range of OD that increases linearly with cell number

The optical density (OD) of a microbial culture increases linearly with the number of cells only for sufficiently small ODs. At higher ODs, the light from the spectrophotometer may scatter off multiple cells, and the relationship between OD and the number of cells becomes non-linear [4]. To calibrate OD measurements, researchers often serially dilute a dense culture of microbes and measure the relationship between the OD and the dilution factor [5, 4] (Fig. 2A). Interpolating this curve, we can convert an OD measurement to the corresponding dilution factor and so correct for any non-linearity between the OD and cell numbers.

**Figure 2.**
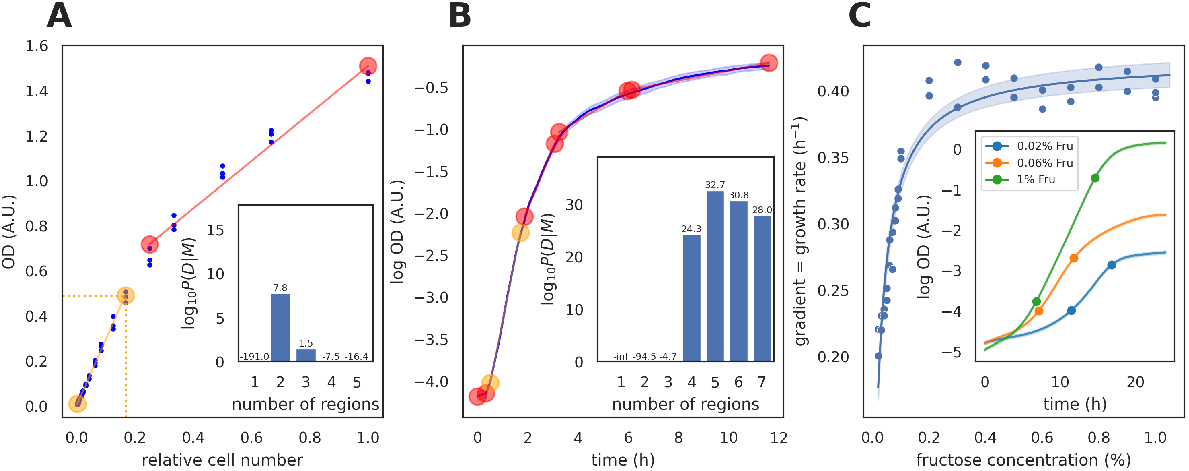
The `nunchaku` algorithm gives intuitive results when applied to biological data. **(A)** The calibration curve for plate-reader measurements of the OD of *Saccharomyces cerevisiae*, found by diluting an overnight culture in 2% fructose, is non-linear (blue dots). There are three replicate measurements for each dilution factor. Our algorithm identifies two linear segments (boundaries marked as circles). Orange circles bound the segment with the highest *R*^2^. We specify the likely maximal range of OD as our prior: [0, 2]. Inset: the logarithm of the model evidence for the number of segments, log_10_ *P*(*D*|*M*). **(B)** Identifying contiguous linear segments in the logarithm of the OD of growing *E. coli* cells as a function of time allows us to identify automatically the region of exponential growth. We show the mean of four replicate measurements (blue) with twice their standard deviation shaded. Circles denote the boundaries of linear segments; orange circles bound the segment with the best-fit line with highest gradient or highest specific growth rate. The average specific growth rate over this segment is 1.5 h^−1^. Inset: log_10_ *P*(*D*|*M*). **(C)** With our algorithm, we can automatically identify the region of exponential growth in multiple data sets, here 38, to reveal growth laws such as Monod’s equation. We plot the specific growth rate in log phase for *S. cerevisiae* as a function of the concentration of fructose, with the solid line a fit of Monod’s equation (*λ*_max_ = 0.423 ± 0.005 h^−1^ and *K*_*M*_ = 0.029 ± 0.002 %). The shaded area shows the 95% confidence interval. Inset: three example growth curves with dots marking the region of exponential growth, identified as the segment with the highest gradient. We specify a prior on the range of *m*, [0, 5].

Dilution factors, however, are not intuitive units, and it is useful to identify the range of ODs over which there is a linear relationship with cell numbers. Not only is this range itself important, but by using the ratio of the maximum of the range to the corresponding dilution factor, we can re-scale the dilution factors back into ODs.

We used the `nunchaku` algorithm to identify the linear range, using the standard deviation over the replicates as the measurement error. Two linear segments are optimal, and the one of interest, where OD is proportional to the number of cells, is the segment beginning at the smallest OD. This segment also has the highest coefficient of determination *R*^2^. Its maximal OD is 0.48 for a relative cell number of 0.17 (Fig. 2A), and we should therefore multiply the dilution factors by 0.48/0.17, or 2.8, to convert back to ODs. Running the algorithm without estimating the measurement error gives a similar result: the conversion factor is 2.76 (Fig. S4A).

### Application 2: Identifying the log phase of microbial growth

Microbes are most often studied when growing exponentially with the log(OD) of the culture linearly increasing with time [2], and researchers identify this log-phase growth from microbial growth curves.

To detect log phase automatically, we applied our algorithm to OD measurements of *Escherichia coli* (Fig. 2B). Partitioning the data into five segments is optimal, and the segment whose best-fit line has the highest gradient — the greatest specific growth rate — corresponds to exponential growth.

Monod noticed an empirical relationship between the nutrient concentration and the specific growth rate of microbes in log phase [2]. Denoting this growth rate as *λ*, the maximal specific growth rate as *λ*_max_, and the nutrient concentration as *s*, his equation becomes

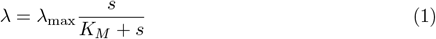

where *K*_*M*_ is now called the Monod constant. To estimate *λ*_max_ and *K*_*M*_, researchers systematically vary the concentration of the carbon source and identify the log phase and the corresponding gradient for each growth curve.

Here we use the `nunchaku` algorithm to provide data to estimate *λ*_max_ and *K*_*M*_ for *S. cerevisiae* growing on fructose (Materials & Methods), from 38 growth curves measured with plate readers (Fig. 2C). Each biological replicate has two technical replicates, and we use a Gaussian process to infer the mean and the standard deviation of the log *OD* from these two replicates [6, 7] (Fig. 2C inset). We use the standard deviation as the measurement error, but re-analysing without this estimate produces similar results (Fig. S4)

## Discussion

Determining where their data is best described by a line is a problem familiar to most scientists. By reframing this question and asking how best to represent a data set as a contiguous series of piece-wise linear segments, we present a statistically rigorous solution.

We use a Bayesian approach, which therefore depends on prior information: the *a priori* bounds on the range of the gradients and intercepts of all possible lines within a segment. The optimal number of segments will depend on this prior if the amount of data is sufficiently small, as it should [8]. In practice, however, users need specify only one range with the other inferred (Materials & Methods), and we see that although a wide prior favours fewer segments, a single segment is robustly assigned to sections of the data that appear linear.

Our method makes two assumptions about how the data deviate from the line segments. We assume these deviations are independent and that each obeys a normal distribution. For some data, a distribution with a purely non-negative support, such as a log normal, may be more appropriate. Although we can use such a distribution in principle, in practice some of the steps that we performed analytically would have to become numerical. If the standard deviation of the normal distribution we use is unknown, we assume additionally that it is identical for all data points. Our algorithm would work too if the standard deviations vary but are proportional to a known function of *x*_*j*_ and *y*_*j*_.

From a wider perspective our work is an example of detecting retrospectively change points in a time series [9]: points where the process generating the dynamics changes. We have simplified this problem by considering change points to occur only at data points and by imposing no continuity on the functions underlying the data for each segment. These simplifications are not restrictive for our task of finding one particular segment of interest. Identifying change points more generally typically requires Markov chain Monte Carlo methods [9, 10], which can call for both more input and expertise from users.

The `nunchaku` algorithm by using enumeration is robust and lends itself to automation, facilitating high throughput studies. It should both ease and increase the reproducibility of data analyses for a wide range of scientists.

## Materials and Methods

### Inferring the most linear regions using model comparison

Given a data set of two variables, we wish to divide the data into the group of contiguous segments that is most piece-wise linear. This problem is different from asking if the data is better described by a linear or a non-linear function. Irrespective of the data’s behaviour, we will find the contiguous segments out of all possible contiguous segments that, piece-wise, are most linear. Our approach answers two questions: how many piece-wise linear contiguous segments best describe the data, and where the optimal boundaries lie for each one.

Let us assume that we have observations, (*x*_*j*_, *y*_*j*_), where *j* runs from 1 to *N* and the *x*_*j*_ are in ascending order. We denote these observations collectively as *D*.

First, we consider whether we should divide the data into *M* or *M*′ segments, using Bayesian model comparison [8]. Assuming equal prior probabilities, *P*(*M*) = *P*(*M*′), we write the Bayes’ factor as:

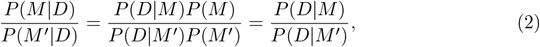

and therefore we should determine the evidence *P*(*D*|*M*) for each *M*.

The evidence is a marginal likelihood. For *M* contiguous segments, there are *M −* 1 unknown boundary points, which we denote as **n** ≡ (*n*_1_, ⋯, *n*_*M−*1_) with *n*_*i*_ *< n*_*i*+1_. The two remaining boundaries are the first and last data points. To fit a line, we need at least three data points, and we assume that each linear segment contains a minimal number of data points *ℓ*_min_, where *ℓ*_min_ = 3, so that *n*_*i*+1_ ≥ *n*_*i*_ + *ℓ*_min_.

The evidence is a sum over all potential **n**:

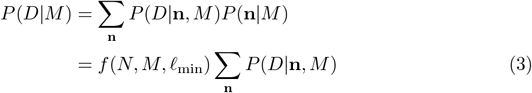

using that any permissible *n*_*i*_ is equally likely to write the prior *P*(**n**|*M*) as a function of *N, M*, and *ℓ*_min_. Specifically, this bounded uniform prior is the reciprocal of the number of possible **n**, which satisfy

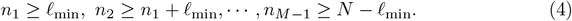

for a given *M* and *ℓ*_min_. We therefore have:

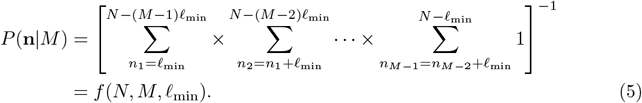

Second, for a given *M* and **n**, we fit the data to *M* lines, each independent of the other. The line ending near the data points indexed by *n*_*i*_ and *n*_*i*+1_ depends only on the data indexed by the indices *n*_*i*_ + 1 and *n*_*i*+1_ inclusively, and this data does not determine any other line. Therefore, mathematically,

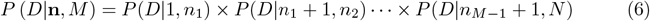

where *P*(*D*|*n*_*i*_ + 1, *n*_*i*+1_) is the likelihood of fitting a line to the data indexed by *n*_*i*_ + 1 to *n*_*i*+1_.

### Finding *P*(*D*|n,*M*)

For each segment of the data, we consider lines with gradient *m* and intercept *c* and let *P*(*y*_*j*_ | *x*_*j*_, *m, c*) describe how data point *y*_*j*_ at *x*_*j*_ deviates from these lines. This deviation is assumed independent of the deviations of other data points. For the *i*’th segment, we then have

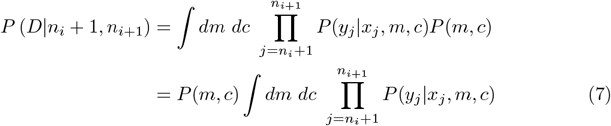

and where we assume that the prior *P*(*m, c*) is a constant, with both *m* and *c* uniformly distributed in some bounded region so that

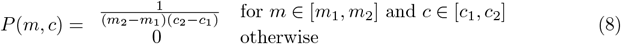

for a fixed *m*_1_, *m*_2_, *c*_1_, and *c*_2_.

### Marginalising *P*(*D*|n, *M*)

Using Eq. 6, we factorise the sum in Eq. 3:

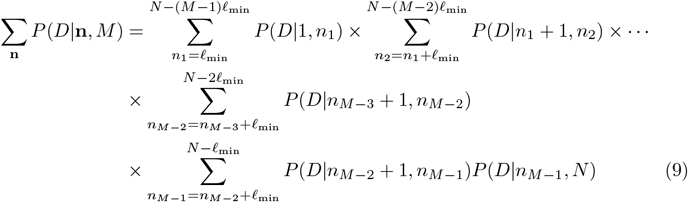

and use the method of variable elimination [11] to evaluate these sums. First we perform the rightmost one, over *n*_*M−*1_, to generate a function of *n*_*M−*2_. We then perform the next rightmost sum, over *n*_*M−*2_, of this function and the next term in Eq. 9, which generates a function of *n*_*M−*3_. We repeat this process until we reach the leftmost sum over *n*_1_, enabling *O*(*MN* ^2^) operations in total instead of *O*(*N* ^*M*^). We evaluate Eq. 5 similarly.

All that remains is to determine *P*(*D*| *n*_*i*_ + 1, *n*_*i*+1_) so that we can find *P*(*D*| *M*) via Eq. 3 and Eq. 9.

### Finding *P*(*D*|*n*_*i*_ + 1, *n*_*i*+1_) for known measurement error

To proceed, we assume that *P*(*y*_*j*_|*x*_*j*_, *m, c*) is a normal distribution with mean *mx*_*j*_ + *c* and a standard deviation *σ*_*j*_. If we know the *σ*_*j*_, for example by approximating each by the corresponding measurement error, then Eq. 7, the likelihood of a line fitting the data indexed by *n*_*i*_ + 1 to *n*_*i*+1_, becomes

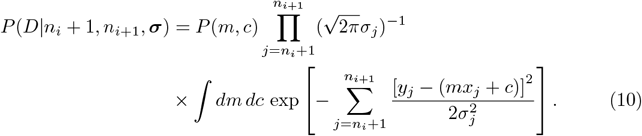

To evaluate the integral, we extend it to infinite range — a suitable approximation because we expect the integrand to be strongly peaked at the most likely values of *m* and *c* [8]. We can then perform the integration analytically. Defining for this segment [12]

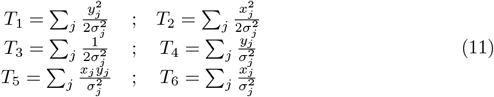

with *j* running from *n*_*i*_ + 1 to *n*_*i*+1_, we find that

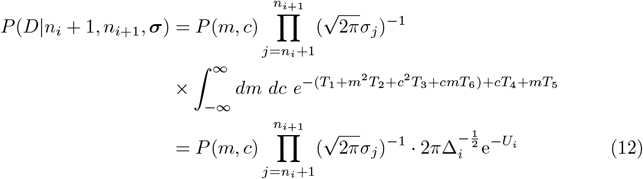

after integrating, where

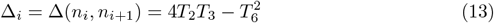

and

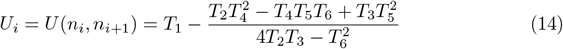

with the subscript indicating the summations in Eq. 11 run from *n*_*i*_ to *n*_*i*+1_.

For this approximation to be valid, we require that the strongly peaked region of (*m, c*) is within the prior range for *m* and *c*. When rewritten as a Gaussian distribution, the integrand of Eq. 12 has a covariance matrix with a determinant Δ_*i*_. The square root of this determinant is proportional to the area under the integrand, and the prior range for *m* and *c* must be large enough to contain this area. Mathematically, using Eq. 8, we need

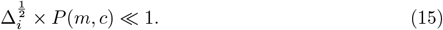

### Finding the boundary points

After determining the optimal number of segments *M*\* into which to divide the data, we next find their boundary points. Using Bayes’ theorem, the posterior for *n* is

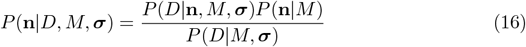

which we evaluate using Eq. 3, Eq. 5, and Eq. 6. We use the mean posterior value of *n*_*i*_ to estimate the optimal *n*_*i*_:

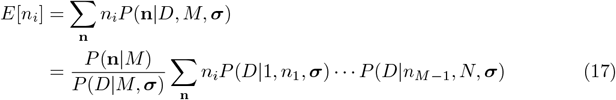

which we sum following Eq. 9. The posterior variance, Var[*n*_*i*_], determines the error in this estimate, which we find similarly.

### Finding *P*(*D*|*M*) for unknown measurement error

If the *σ*_*j*_ are unknown, we assume the same constant *σ* for all *j* with a uniform prior probability between [*σ*_min_, *σ*_max_] [13]. Eq. 3 then becomes

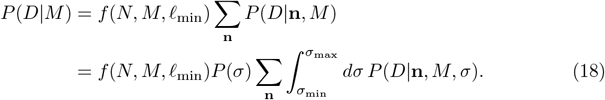

The constant *P*(*σ*) = 1/(*σ*_max_ *− σ*_min_) will cancel in Eq. 2 when we compare the evidence for different *M*.

To evaluate the integral, we write the number of data points contributing to the *i*’th linear segment as *ℓ*_*i*_ = *n*_*i*+1_ *−n*_*i*_, and re-define the *T*_1_ to *T*_6_ in Eq. 11 by removing their dependence on *σ* so that

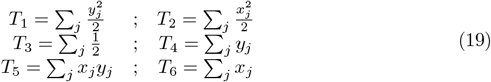

with *j* running from *n*_*i*_ + 1 to *n*_*i*+1_. Then, similar to Eq. 12,

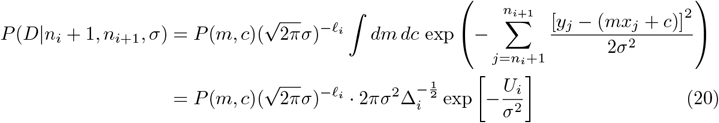

with Δ_*i*_ = Δ(*n*_*i*_, *n*_*i*+1_) and *U*_*i*_ = *U*(*n*_*i*_, *n*_*i*+1_) defined in Eq. 14, with Eq. 19. Consequently,

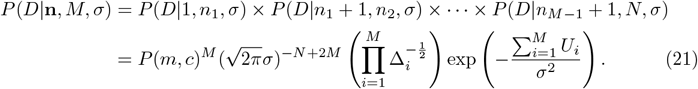

Although with Eq. 21 it is possible to approximate analytically the integral over *σ* in Eq. 18 by extending the range of the integrand to (0, *∞*), the resulting expression prevents us from summing over **n** using variable elimination. Instead, we write

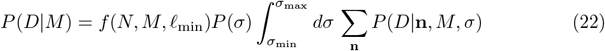

and numerically evaluate the integral, using variable elimination to sum over **n** in Eq. 22 for each *σ* chosen by the integration algorithm. We find the expected boundary points via Eq. 17, again numerically integrating over *σ*.

### Performing the integration

To stabilise the numerical integration, we scale the integrand of Eq. 22 by its value at the most likely value of *σ*, making the integrand nearly always less than one and preventing overflow. We use expectation-maximisation to estimate the most likely *σ* for a given *M*. The EM algorithm finds the *σ* that maximises *P*(*D*|*M, σ*) [14]. We guess a value of *σ, σ*_*o*_ say, and find *P*(**n**|*D, σ*_*o*_, *M*) from Eq. 16. To update *σ*_*o*_, we maximise *Q*(*σ, σ*_*o*_) with respect to *σ*, where

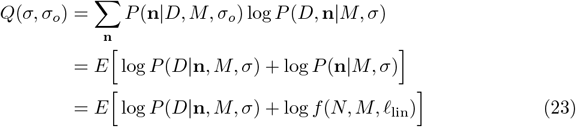

with the expectations taken over *P*(**n**|*D, M, σ*_*o*_). Expanding Eq. 23 using Eq. 21, there are only two terms that depend on *σ*, and we can differentiate to find the updated *σ* = *σ*_*n*_:

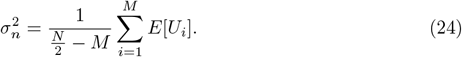

We use the equivalent of Eq. 17 with ***σ*** = *σ*_*o*_ to evaluate these expectations and iterate until the value of *σ* converges.

### Implementation

To compare the different linear segments, we calculate the gradient, intercept, and the coefficient of determination *R*^2^ of the line maximising the likelihood for each. The user can then select a desired segment, such as the one with the largest gradient.

The algorithm requires the *a priori* ranges of *m* and *c*: [*m*_1_, *m*_2_] and [*c*_1_, *c*_2_] in Eq. 8.

The user can either provide both ranges or only [*m*_1_, *m*_2_] or the maximal range of *y* possible in the experiment, [*y*_min_, *y*_max_].

If the user provides only [*m*_1_, *m*_2_], we estimate *c*_1_ as min (*−m*_2_*x*_max_, *m*_1_*x*_min_) and *c*_2_ as max (*−m*_1_*x*_max_, *m*_2_*x*_min_).

If the user provides the range of *y*, we estimate [*m*_1_, *m*_2_] as [*−m*_max_, *m*_max_], with *m*_max_ = (*y*_max_ *− y*_min_)/Δ*x*_min_ and Δ*x*_min_ being the smallest difference between two neighbouring *x* values.

### Availability

We coded the algorithm as a Python package, `nunchaku`, available at https://pypi.org/project/nunchaku and via pip. We have also embedded `nunchaku` into our omniplate software for analysing plate-reader data [7].

### Generating and testing with synthetic data

To test our method, we generated a piece-wise linear function *f*(*x*) with 1 *≤ M ≤* 10 continuous linear segments, each having between 10–50 data points and with a unit distance, Δ*x* = 1, between data points. We sampled *θ*, the angle between each segment and the *x*-axis, from a uniform distribution on the interval [−tan^−1^(20), tan^−1^(20)], so that the gradient, tan *θ*, lies between [−20, 20]. Furthermore we ensured that the difference in *θ* between neighbouring segments is larger than a fixed minimum, *θ*_0_. We added Gaussian noise *ϵ ∼* Normal(0, *σ*^2^) to give 10 replicates of *y* = *f*(*x*) + *ϵ*. We generated 3,600 synthetic data sets in total, a combination of 200 different piece-wise linear functions *f*(*x*), three values of *θ*_0_, and six values of *σ*^2^. In Fig. 1, *θ*_0_ = 10°.

## Experimental methods

We used a prototrophic strain of *S. cerevisiae* (FY4), pre-cultured in synthetic complete (SC) medium with 2% (w/v) sodium pyruvate in a 30°C shaking incubator at 180 rpm for two days. Before the experiment, we diluted the cells six-fold and let them grow for six hours. After washing the cells twice with fresh minimal media [15], we inoculated them into minimal media with different concentrations of fructose on a 96-well microplate. The liquid volume of each well was 200 *µ*l.

For *E. coli*, we pre-cultured cells in 3 ml liquid Luria broth (LB) with one colony from a fresh plate and grew aerobically to log phase (6h) at 37°C with 250 rpm shaking. We then inoculated 3 *µ*l culture into 147 *µ*l fresh LB medium per well on a 96-well microplate.

We used either a Tecan Infinite M200 Pro or F200 plate reader at 30°C for *S. cerevisiae* and 37°C for *E. coli* with linear shaking at amplitude 6 mm. Measurements of absorbance at 600 nm, OD_600_, were taken every 10 minutes.

### Fitting Monod’s equation

After estimating the specific growth rate *λ* at each concentration of fructose *s*, we have a data set *D ≡ {*(*λ*_*i*_, *s*_*i*_)*}* with 38 data points. We use Bayesian inference to estimate the constants *λ*_*K*_ and *K*_*s*_ of Monod’s equation. Assuming a Gaussian measurement error of *λ*_*K*_ with a standard deviation *σ* and independent measurements, the likelihood

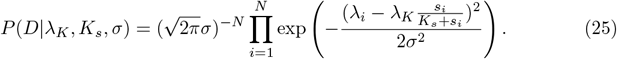

To marginalise over *σ*, we assume *P*(*σ*) ∝ 1/*σ*, so that

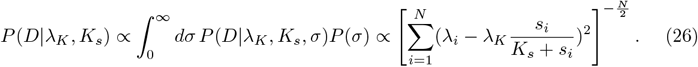

We further assume that the prior *P*(*λ*_*K*_, *K*_*s*_) is uniform, making the posterior probability *P*(*λ*_*K*_, *K*_*s*_|*D*) proportional to the likelihood. We therefore maximise the likelihood with respect to *λ*_*K*_ and *K*_*s*_ using the Broyden–Fletcher–Goldfarb–Shanno (BFGS) algorithm. We estimate the errors in these inferences using the diagonal elements of the Hessian matrix −∇∇ log *P*(*D*|*λ*_*K*_, *K*_*s*_) evaluated at the maximum of the likelihood [16].

## Acknowledgements

We thank Ramon Grima and Edward WJ Wallace for helpful comments and the BBSRC (PSS & YH) and the Darwin Trust (YH) for funding.

This research was funded in whole, or in part, by the BBSRC [BB/W006545/1]. For the purpose of open access, the authors have applied a creative commons attribution (CC BY) licence to any author accepted manuscript version arising.

## Supplementary Figures

**Figure S1.**
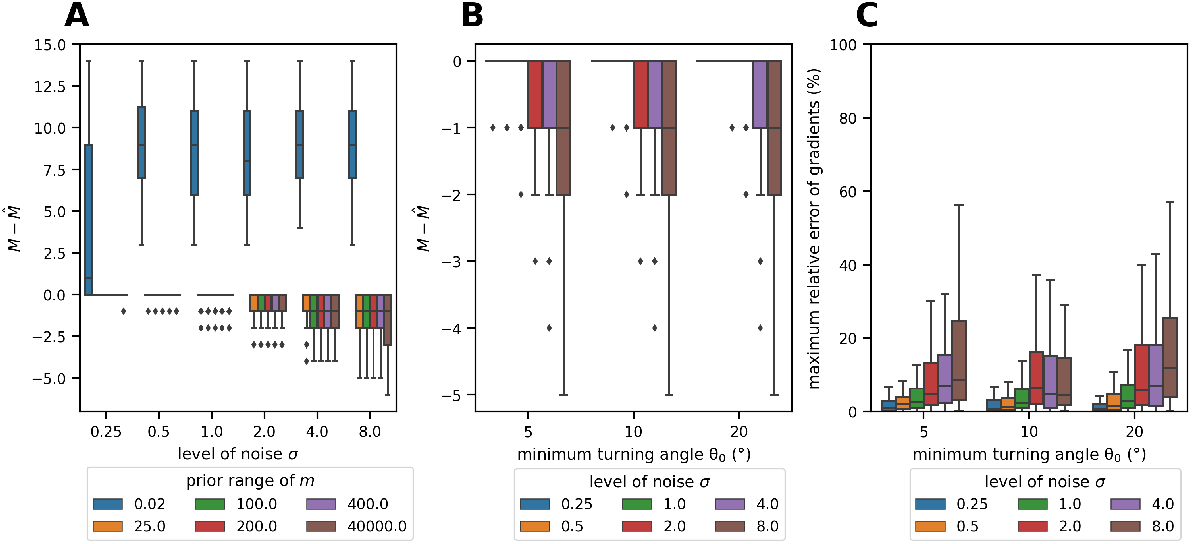
The performance of `nunchaku` on synthetic data. **(A, B)** We show how the difference between the true number of linear regions *M* and the number estimated by Nunchaku 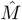 depends on (A) the level of noise *σ* and the *a priori* range of the gradient and (B) the minimum angular difference between neighbouring segments *θ*_0_ and the level of noise *σ*. We assume a symmetric prior: for example, an *a priori* range of 25 means [−25, 25]. **(C)** The maximum relative error in the estimated gradients depends on the complexity of the problem, which we specify with *θ*_0_ and *σ*. We calculate this error only when 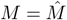. For each function *f*(*x*) with *M* linear segments and gradients (*m*_1_, ⋯, *m*_*M*_), we estimate the relative percentage error of the estimated gradient 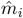 for each segment *i*: 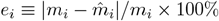, and we plot max_*i*_ *e*_*i*_. The *a priori* range of the gradient is [−25, 25], and we generated the 3,600 data sets following Methods.

**Figure S2.**
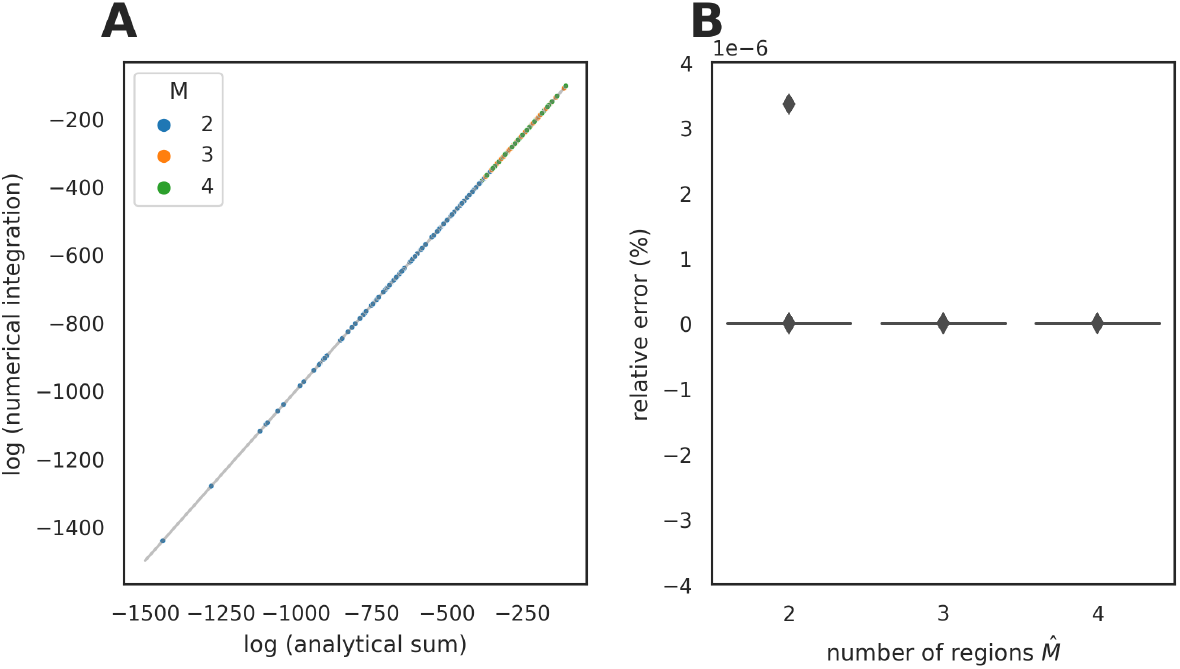
Validating the numerical integration when the measurement error is unknown. **(A)** We plot Eq. 21 analytically integrated over *σ* and then summed numerically over **n** against Eq. 22 with its integral evaluated numerically. The gray line shows *y* = *x*. **(B)** Box plots of the percentage error of the numerical integration relative to the analytical sum at three different values of 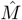 when *M* = 3 show that the error is negligible. We generated 200 data sets from 200 different piece-wise linear functions with *σ* = 1, *θ*_0_ = 10°, and *M* = 3. Each data set has two replicates of *y*.

**Figure S3.**
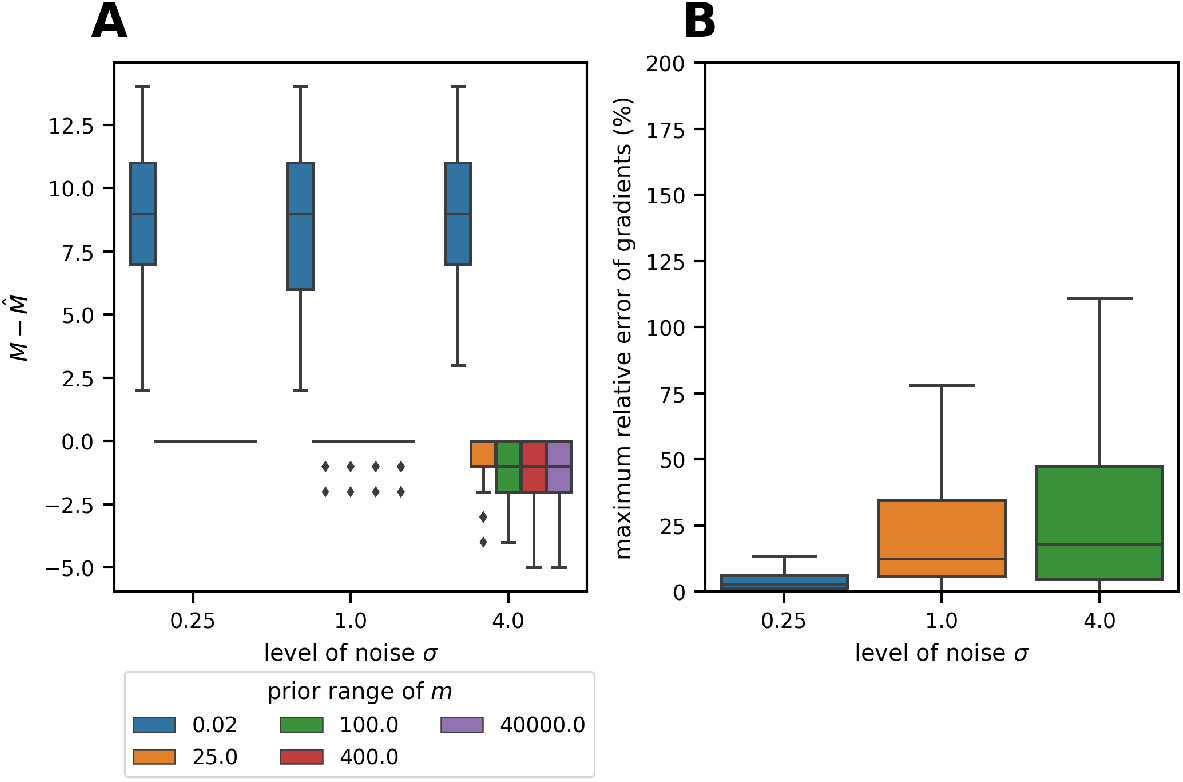
The performance of `nunchaku` on synthetic data when the measurement error is neither provided nor estimated. **(A)** We show how the difference between the true number of linear regions *M* and the number estimated 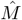 depends on the level of noise *σ* and the *a priori* range of the gradient. Performance is worse than in Fig. S1A. **(C)** The maximum relative error of the estimated gradients as a function of *σ* is larger than Fig. S1C. We generated 800 data sets, with a combination of 200 different piece-wise linear functions, three values of *σ*, and *θ*_0_ = 10°. Each data set has one replicate of *y*.

**Figure S4.**
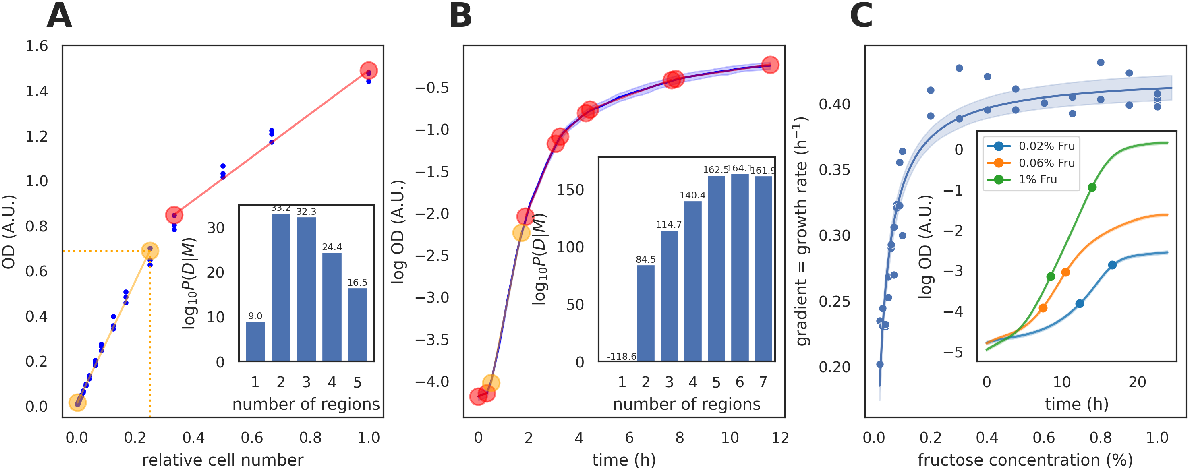
The `nunchaku` algorithm gives similar results to Fig. 2 when the measurement error is neither provided nor estimated. **(A)** We plot the OD a function of the cell number relative to an overnight culture of *S. cerevisiae* in 2% fructose (blue dots) and the linear regions identified with nunchcaku. The orange section ends with the data point adjacent to the one chosen in Fig. 2. Inset: log_10_ *P*(*D*|*M*). **(B)** The logarithm of the OD of a culture of *E. coli* plotted as a function of time. Inset: log_10_ *P*(*D* |*M*). The blue curve shows the mean log(OD) of four replicates with the shading twice the standard deviation. The red circles in (A) and (B) mark the end points of each identified linear region. We find each red line using linear regression on the data for that segment. Orange highlights the region of interest: the region with the largest *R*^2^ in (A) and the region with the largest gradient in (B). **(C)** The specific growth rate, the gradient of the log(OD), for *S. cerevisiae* growing in log phase as a function of fructose concentration. The solid line shows a fit of Monod’s equation, Eq. 1, with *λ*_max_ = 0.422 ± 0.006 h^−1^ and *K*_*M*_ = 0.026 ± 0.002%. The shading shows the 95% confidence interval. Inset: three example growth curves with the dots on each curve marking the end points of log phase identified by `nunchaku`. These data are noisy for some concentrations of fructose. We then ignored the first hour of measurements where the signal-to-noise ratio is particularly poor and increased *ℓ*_min_ to five. We used [0, 2] as the prior range of *y* in (A) and [0, 5] as the prior range of *m* in (B) and (C).

## Notes

### Competing Interest Statement

The authors have declared no competing interest.

